# Navigators learn a local graph, not a global map

**DOI:** 10.64898/2026.01.11.698858

**Authors:** Marianne Strickrodt, Tobias Meilinger, Heinrich H. Bülthoff, William H. Warren

**Affiliations:** Max-Planck-Institute for Biological Cybernetics, Max-Planck-Ring 8-14 72076 Tübingen, Germany; Graduate Training Centre of Neuroscience, Österbergstraße 3, 72074 Tübingen, Germany; Applied Cognitive Psychology, University of Tübingen, Schleichstr. 4, 72076 Tübingen, Germany; Department of Cognitive, Linguistic & Psychological Sciences, Brown University, 190 Thayer Street, Providence, RI 02912, USA

**Keywords:** navigation, spatial memory, cognitive map, Euclidean map, graph knowledge, labeled graph, network of reference frames, impossible environments, virtual reality

## Abstract

It is still an unresolved issue how humans represent navigable space and use this information to relate distant locations, as when shortcuting or pointing. In the present study we compare two competing theoretical approaches to the structure of this spatial knowledge. We contrast the metric embedding of spatial information in a global reference frame (i.e., a Euclidean mental map) with models that assume only local place-to-place information (i.e., a labeled graph), which can be used to estimate shortcuts and pointing direction when needed. Two groups of participants learned a multi-corridor virtual maze by walking around a zig-zag loop that connected seven objects placed on a circle. One group experienced a possible Euclidean maze, and the other group an impossible, non-Euclidean, ‘broken’ version of the maze. In the impossible environment, after walking one lap the participant was covertly teleported to the starting place again, despite having walked to a different Euclidean location. Thus, the local place-to-place metrics were globally inconsistent. During the test phase, participants pointed to targets in a clockwise or counterclockwise sequence around the circle from their current location. Whereas the possible maze group was fairly accurate, the estimates of the impossible maze group were systematically biased by the test sequence, as predicted by local place-to-place metrics. Despite being queried about the same target, participants pointed in significantly different directions, violating the metric postulates. The results suggest that human knowledge of navigable space is not a globally consistent Euclidean map, but can be characterized as a labeled graph.

## Introduction

As we navigate the world every day, we travel to many different places. When going grocery shopping, we leave our apartment, walk a few blocks, passing our hair dresser on the way to the supermarket. On our way back, we might take a detour to stop by a friend’s house and head home afterwards. A characteristic of such navigable environments is that they cannot be viewed from a single vantage point but must be experienced successively. Nonetheless we are able to apprehend the relative locations of the places visited and form survey knowledge of the neighborhood, which allows us to approximate straight-line directions and distances and take novel shortcuts across unexplored terrain between learned places.

There are a range of hypotheses about the structure of such survey knowledge. One influential proposition is the *cognitive map*. As formulated by Tolman (1948), it was “something like a field map of the environment… indicating routes and paths and environmental relationships” (p. 192). Even though Tolman may not have intended his “cognitive map” to be interpreted literally, the term was adopted and expanded by other researchers in this vein (e.g., Byrne, Becker, & Burgess, 2007; Gallistel, 1990; O’Keefe & Nadel, 1978; Siegel & White, 1975). On this interpretation, a cognitive map is a metric Euclidean mental representation of environmental space. The discovery of (among others) place cells and grid cells in rats (e.g., Hafting, Fyhn, Molden, Moser, & Moser, 2005; O’Keefe & Nadel, 1978) and later place and grid-like activity in humans (e.g., Doeller, Barry, & Burgess, 2010; Ekstrom et al., 2003; Jacobs et al., 2013; Jacobs, Kahana, Ekstrom, Mollison, & Fried, 2010) suggests a neural basis for the embedding of place information into a stable, metric reference system. A key component of the cognitive map approach is that all the places we experience can be assigned distinct locations in the cognitive reference system, similar to (x, y) coordinates in a Cartesian coordinate system (e.g., Gallistel, 1990; Gallistel & Cramer, 1996). In such a format “every point on the map is related to every other point within the reference frame of the map” (Nadel, 2013, p. 166). Survey tasks such as distance and direction estimation to non-visible landmarks or novel shortcuts can be based on such a cognitive map by reading out coordinates and calculating distances and angles between them.

An example is represented in Figure 1, where four places (J, K, L and M), which are experienced successively in a circular manner in the external world (panel a), are represented in the form of a Euclidean map (panel b). Such a global Euclidean cognitive map must satisfy the metric postulates of positivity, symmetry, and triangle inequality (e.g., Beals, Krantz, & Tversky, 1968; McNamara & Diwadkar, 1997). Positivity refers to the idea that the distance between any point and itself must be zero, as there can only be one point in the Euclidean mental map representing a place in the external world. Further, the distance between any two points must be larger than zero, as representation of distinct places cannot overlap. Symmetry is achieved when the distance estimated from point A to B is the same as the distance estimated from point B to A, such that the vector from A to B is the exact reverse from B to A. The triangle inequality defines the relationship between any three points such that the sum of the distances between points A and B and between points B and C must always be larger or equal to the distance between A and C. Further, for a Euclidean metric the inner angles of such a triangle should sum up to 180°. Similarly, the four points in panel B must form an irregular quadrilateral with a sum of inner angles of 360°.

**Figure 1.**
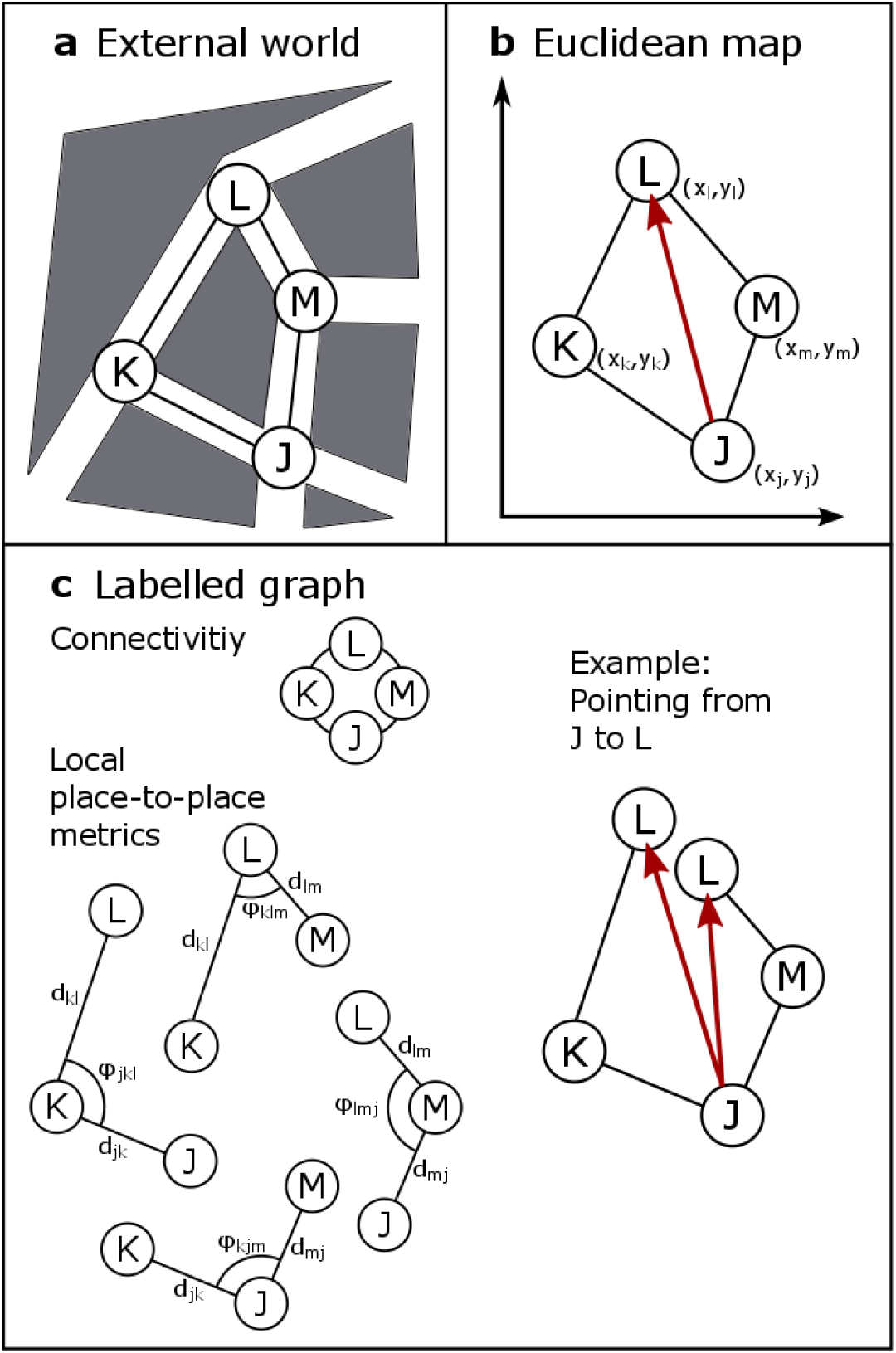
Contrasting predictions for Euclidean map and labeled graph representation. **(a)** Example of navigable space, four places within a street network (J, K, L M) connected successively. **(b)** Distorted but globally consistent Euclidean map representation of panel a. Each place is assigned to a distinct coordinate which can be read-out from the coordinate system to make survey estimates. **(c)** Labeled graph representation consisting of connectivity and local metrics. Note that, albeit redundant, distances are visualized twice for clarification reasons. Translational and rotational metrics are restricted to the direct neighbors of a place and need not to be globally consistent, leading to violation of metric postulates when used for survey estimates of distant places. For example, recalling the location of place L when standing at place J: depending on which local place-to-place metrics are used (clockwise or counterclockwise) different estimates of the location of place L are possible.

A Euclidean mental map need not be a perfect representation of the external world, however. We know, for example, that the human path integration suffers from a rather low resolution and discontinuities (e.g., Loomis et al., 1993; Zhao & Warren, 2015a, 2015b), and that boundaries bias distance estimates between places (e.g., Kosslyn, Pick, & Fariello, 1974; McNamara, 1986; Newcombe & Liben, 1982). Thus, erroneous and distorted representations are likely, but can still satisfy the metric postulates.

Several findings seem to challenge the Euclidean map hypothesis. Yet most of them can be explained by additional assumptions, levels of representation, recall mechanisms or map distortions. For instance, humans tend to remember irregular environments and junctions as more regular (e.g., orthogonal streets) than they are (e.g., Byrne, 1979; Moar & Bower, 1983; Tversky, 1981). However, participants are still able to draw a coherent map, suggesting a distorted Euclidean representation (e.g., Tversky, 1981). Moar and Bower (1983), who obtained angular judgments between triplets of interconnected intersections, noted that most of the intersections they tested were indeed orthogonal, and the non-orthogonal relation between the triplets only became apparent when following curvy streets connecting two intersections. The salient memory of orthogonal intersections might have been used for the angular judgments between intersections instead of the actual survey relations stored in memory, for example, in the form of a Euclidean map.

Other findings suggest a contradiction of the metric postulate of symmetry for Euclidean mental maps. Spatial judgments between the same pair of places differ depending on whether they are made from place A to B or from place B to A. Such asymmetries were found in route selection (Stern & Leiser, 2010) and distance estimations (Burroughs & Sadalla, 1979; McNamara & Diwadkar, 1997; Sadalla, Burroughs, & Staplin, 1980). However, two models have been proposed to account for such asymmetries while leaving the assumption of a Euclidean mental map with symmetric distance representation untouched. Both the category-adjustment model of spatial coding (Huttenlocher, Hedges, & Duncan, 1991; Newcombe, Huttenlocher, Sandberg, Lie, & Johnson, 1999) and the contextual-scaling model (McNamara & Diwadkar, 1997) locate the source of asymmetries in the estimation process, either by assuming an additional layer of representation (category layer) or more or less salient memory content unique to each reference object, which may interact with the Euclidean map layer. Both models render the observed asymmetries much less of a violation of metric postulates than originally thought.

The character of survey knowledge has also been probed using impossible, non-Euclidean environments. The underlying assumption of these studies can be stated succinctly: If a Euclidean map is the underlying format of human spatial knowledge, experience in impossible environments should be embedded into a geometrically consistent format and survey tasks solved accordingly. Kluss, Marsh, Zetzsche and Schill (2015) had participants walk impossible virtual environments on a treadmill while wearing a head-mounted display and subsequently reproduce one walk through each environment blindfolded. For example, in one impossible environment three corridors formed a triangle, but were connected by 90° angles (the sum of inner angles equaled 270° instead of 180° as in the possible environment). Turning angles during reproduction closely resembled the experienced angles (sum of angles around 270°), indicating that local metric but globally inconsistent knowledge was acquired. However, due to high variability in responses, the sum of angles in the impossible and possible triangles did not differ significantly, so a global Euclidean embedding could not be rejected.

In a study by Warren, Rothman, Schnapp, & Ericson (2017), participants learned the location of eight objects within a complex, 11m x 11m virtual hedge maze by walking in a large tracking hall while wearing a head-mounted display. For one group of subjects, the maze contained two virtual “wormholes” that teleported the participant to another maze location, rotated by 90°, unbeknownst to them. When participants were instructed to perform straight line shortcuts between the learned objects, strong biases of 37° towards the wormhole locations (out of the expected 45°) were found for targets near wormhole entrances. In particular, the same object was remembered in two physical locations 6m apart, thus indicating a violation of the positivity postulate in spatial knowledge. A second experiment found that near-wormhole locations were remembered as ripped away from close-by objects and folded over the locations of other objects. This result not only violated the positivity postulate but also indicated ordinal reversals in remembered spatial locations. Indeed, using a 3D wormhole maze Murvy and Glennerster (2018) found that pointing behavior could be best modelled by assuming local distortions in the locations and orientations of places rather than a geometrically consistent map. In sum, these studies show that the local metrics experienced from one place to the next strongly influence the structure of spatial knowledge. They imply that human survey knowledge is not best characterized as a metric Euclidean map.

Importantly, there is no need for a global metric embedding of places to perform survey tasks. Alternatively, performance could be explained by spatial knowledge about the connectivity of places, such as a topological graph in which nodes correspond to places and edges to paths linking them, augmented by local metric information about the rotation and translation between places (e.g., Chrastil & Warren, 2014; Meilinger, 2008; Warren, Rothman, Schnapp, & Ericson, 2017). Albeit differing in detail, such theories can be summarized under the term *labeled graph theories*. Figure 1c visualizes the idea. For example, when learning the connectivity of J-K-L we additionally acquire the approximate distance between K and M as well as that between K and J. Furthermore, approximate angular information about the direction of J relative to L when standing at K is acquired. Such local place-to-place metrics are learned for the other neighboring places as well.

To perform a survey task, for example standing at J and pointing to L, one’s current position and the target must be brought into direct reference by successively recalling the local metrics from one’s current position to the target to make an estimate. This can be done by vector addition (Warren et al., 2017), for example by imagining how the nonvisible parts of the environment could be strung together successively (Meilinger, 2008). Like the Euclidean map, the labeled graph incorporates errors in those local metrics that arise during the encoding process. The crucial difference between the two approaches is that in a labeled graph these local measurements are not embedded into a common coordinate system, and do not need to be globally consistent.

An example appears in Figure 1c. Suppose the navigator is estimating the direction from J to L, based on the local graph information. Performing vector addition clockwise, from J to K to L, results in a different estimate from doing so counterclockwise, from J to M to L, because the local metrics in the two directions are inconsistent. Such distortions might remain unnoticed in the majority of studies due to the very high absolute angular errors of 20-100° observed in such pointing tasks (e.g., Chrastil & Warren, 2013; Foo, Warren, Duchon, & Tarr, 2005; Ishikawa & Montello, 2006; Meilinger, Riecke, & Bülthoff, 2014; Schinazi, Nardi, Newcombe, Shipley, & Epstein, 2013; Weisberg, Schinazi, Newcombe, Shipley, & Epstein, 2014). As a consequence of errors in local place-to-place metrics, the metric postulates of positivity and triangle inequality can be violated on the global scale. There is not one distinct location for each place and calculating the sum of the inner angles between the four places shown in our scenario in Figure 1 does not necessarily correspond to 360°, as expected for a Euclidean metric.

Not only is the labeled graph hypothesis able to explain previous findings that are inconsistent with a Euclidean map, but it can also account for a number of additional findings. In a recent study, Ericson and Warren (2020) investigated the invariant properties of spatial knowledge by subjecting a virtual hedge maze to various transformations during learning, and found results supporting the labeled graph hypothesis. There is ample evidence for the formation of distinct memory units for individual local environments. Direction estimation in a multi-corridor space is facilitated when the body is aligned with the view first experienced within each corridor (i.e., when looking along the corridor), compared to being mis-aligned (e.g., facing the wall) (e.g., Meilinger et al., 2014; Strickrodt, Bülthoff, & Meilinger, 2018). This is typically interpreted as the formation of a local reference frame confined to the visible environment or ‘vista’. Locations of objects within a vista are stored relative to this local reference frame. Being aligned with the reference axis of the mental reference system facilitates the recall of positional information, while being mis-aligned involves effortful mental transformations to map the orientation of the reference system onto one’s current view (e.g., McNamara, Sluzenski, & Rump, 2008; Mou, McNamara, Valiquette, & Rump, 2004).

In addition, pointing latencies increase with the number of traversed corridors between the current location and the target (e.g., Meilinger, Strickrodt, & Bülthoff, 2016; Meilinger, Strickrodt, & Bülthoff, 2018; Strickrodt et al., 2018). Such corridor distance effects seem to reflect the recall process expected for vector addition through a labeled graph, corresponding to the successive activation of remembered places along a path through the graph. Furthermore, knowing the exact location of an object within a vista space does not seem to be accompanied by knowledge about where the vista space itself is located (Marchette, Ryan, & Epstein, 2017). Such findings are in accordance with the labeled graph hypothesis, where places are connected by edges forming a graph. At the same time they do not disprove the Euclidean map hypothesis, since graph information could be embedded in an additional layer. Distance effects in pointing latency could originate from the graph layer utilized also in direction estimates.

The aim of the present study was to compare specific predictions of the Euclidean map and labeled graph hypotheses by systematically biasing the local metric information in an impossible non-Euclidean environment. Given that a labeled graph encodes local place-to-place metrics, inconsistent metric information should be stored as it is experienced. A Euclidean map, however, embeds local measurements in a global coordinate system, so when faced with inconsistent local metrics, local adjustments are made to regularize the measurements and achieve a globally consistent embedding (see e.g., Mallot & Basten, 2009; Wang, 2016). We thus designed a virtual environment that would make different predictions based on local place-to-place metrics and a globally consistent embedding

The present study thus tests two models of human survey knowledge by contrasting quantitative predictions for reliance on either local place-to-place metrics (labeled graph) or a global metric embedding (Euclidean map). One group of participants learned a possible multi-corridor virtual maze by walking around a zig-zag loop that connected seven objects placed in a circle (Figure 2a). The other group learned an impossible, non-Euclidean, ‘broken’ version of the maze (Figure 2b), in which object locations were shifted and participants were covertly teleported across a physical distance gap between virtually adjacent corridors. In the test phase, they were then asked to estimate the straight-line direction between pairs of objects by pointing. Critically, target objects were tested in either a clockwise or counterclockwise sequence around the ring. If a labeled graph underlies the estimation process, pointing estimates should reflect the local place-to-place metrics that are activated by the test sequence. The graph hypothesis thus predicts strongly biased responses to targets in the impossible maze compared to the possible maze. In particular, pointing directions to the same targets should differ when tested in the clockwise and counterclockwise sequence, thereby violating the positivity postulate (compare Figure 1c). In contrast, if the local information is globally embedded, the place-to-place metrics around the circle of objects should be regularized to form a geometrically consistent map that satisfies the metric postulates. The map hypothesis thus predicts that pointing directions should be similar when tested in the clockwise and counterclockwise sequence, in both the possible and impossible environments.

**Figure 2.**
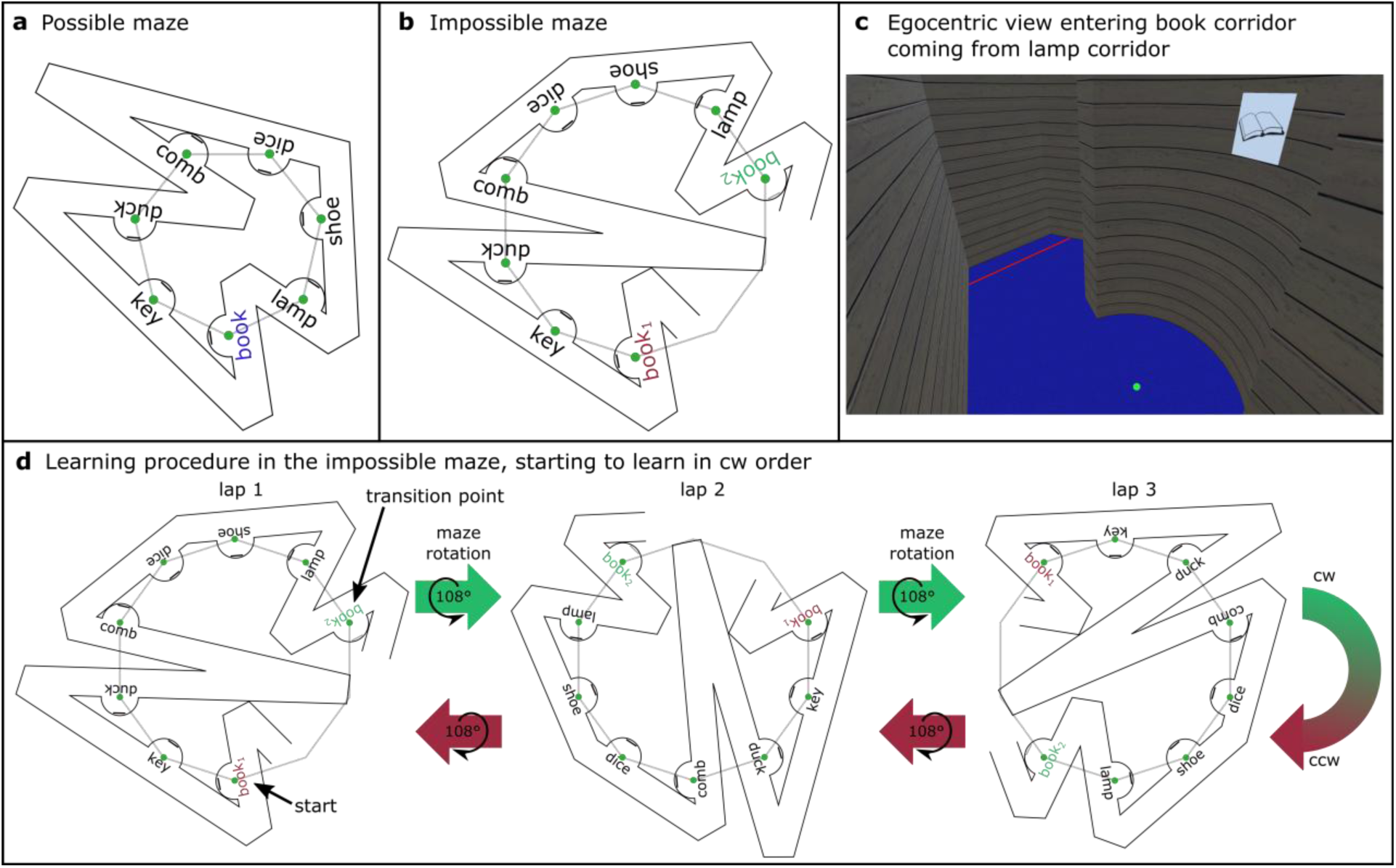
Design of the virtual environments. **(a)** Possible maze. Seven corridors are connected to form a zig-zag loop with left and right turns. Seven places (marked by green dots) are evenly spaced along the vertices of a heptagon (grey). Hence, global and local metrics match up to form a Euclidean space. **(b)** Layout of the impossible maze, a widened-up version of the possible maze. Seven places (green dots) are spaced at seven vertices of a decagon (grey), forming a non-Euclidean world. Two identical book corridors (book_1_ and book_2_) allow for unnoticeable teleportation. **(c)** Example of an egocentric view. A green dot on the floor indicated the exact place that had to be learned and was identified by the object hanging in the center of the alcove wall. A wooden panel structure was attached to the walls, a blue carped on the floor. **(d)** Example of learning procedure when starting clockwise (cw). Lap 1 (left) is started at the book corridor (red book_1_) and continued clockwise (key, duck, etc.), until reaching a book corridor identical to the first one (green book_2_) without having reached the original location in physical space. At this point the virtual maze was rotated by 108° (middle) to match up the first book corridor (red book_1_) with participants current position. Hence, visually seamless lap 2 started and participants continued walking cw till reaching the book corridor distant from where they started off again (green book_2_). The maze was rotated again (right). When reaching the book corridor a third time after lap 3 participants changed walking direction and returned along their previous path in the counterclockwise corridor order for another three laps (panel from right to left). Like this the local corridor-to-corridor metrics remained constant across each lap although the space was not matching up on a global scale in physical space.

## Method

### Participants

Forty-eight participants with normal or corrected to normal visual acuity took part in the experiment, all naïve to the purpose of the research, and received monetary compensation for their participation. Twenty-five of them were assigned to the possible maze, and 23 to learn the impossible maze. Two participants failed to complete the experiment, one because the task was too hard and the other due to equipment malfunction; another ten participants were excluded from analysis due to a chance level performance in the testing phase, six from the possible maze group and four from the impossible maze group (see Results). This left 18 participants in the possible group (M 7, F 11, age *M*=29.61, *SD*=9.73) and 18 participants in the impossible group (M 5, F 13, age *M*=28.94, *SD*=12.46), representing competent navigators in the top three quartiles of participants. This sample size was based on results from a preliminary study for the present experiment. The experiment was approved by the local ethics committee.

### Material

Participants walked freely in a large tracking area (12×12 m) while viewing the virtual environment in a head-mounted display (Oculus CV1 HMD) with a 110° diagonal field of view and a resolution of 1080 x1200 pixels per eye. Head position was recorded with 20 high-speed infrared cameras (Vicon® MX 13) at 180 Hz and transmitted wirelessly to a backpack computer (MSI VR One, with NVIDIA GTX 1070 graphics) carried by the participant. This immersive virtual reality setup provided both visual and idiothetic information about self-motion. Bird sounds were played via the HMD’s headphones to mask any directional auditory cues. The experimenter followed the participant closely to ensure their safety and to eliminate fixed auditory cues when giving instructions.

The virtual environment consisted of seven connected corridors, varying in length and connecting angle, that formed a zig-zag loop with a total length of 33.35m (see Figure 2a and b). After passing through the seventh corridor participants would end up back in the first corridor again. This loop structure enabled participants to continuously walk through the environment for multiple laps (clockwise or counterclockwise). Each corridor (90cm wide) contained a semi-circular alcove (60 cm diameter) on the inner wall, with a picture of an object centered on the wall at about eye height (Figure 2c). The alcoves appeared on the inner wall of the corridors so they would always be on the participant’s right when walking clockwise around the loop, and vice versa.

The center of each alcove was marked by a green dot on the floor, indicating the places participants had to learn. Thus, each place was unambiguously localized by the green dot and labeled by the object in the alcove (Figure 2c). The objects were black-and-white line drawings on a 25 x 25cm plane (2D) with monosyllabic names: book, key, duck, comb, dice, shoe, lamp (clockwise order). The experiment was conducted in English. The identifiability of the drawings was verified in a pretest, in which four additional participants, three of them native English speakers, named all items correctly and unambiguously.

In the possible maze (Figure 2a), the seven places (green dots) were evenly spaced around a circle, at the vertices of a symmetric heptagon (i.e., seven-sided polygon). The side length of the heptagon was 2.38m, corresponding to straight-line distance between adjacent neighbors (e.g., dice and shoe), and the interior angle between adjacent sides of the heptagon was 128.57°. All places were evenly spaced around the circle, and when standing at one place there was an angular difference of 25.71° between any pair of adjacent objects (e.g., standing at the dice, and pointing to the shoe and lamp). Because of the symmetry of the heptagon, the same holds true for all seven pointing positions, allowing us to average the responses across positions for analysis. Throughout the experiment, the visual gain was matched to the participants physical head movements.

The impossible maze (Figure 2b) was a distorted version of the possible maze. We broke the connection between the first corridor (book) and the seventh corridor (lamp) and pulled the maze apart, so the seven places were relocated on a larger circle, corresponding to adjacent vertices of a decagon (ten-sided polygon), leaving a large gap of three vertices. The side length of the decagon was 2.38m, the same as the possible maze, but its interior angle was 144°, and when standing at one place the angular difference between any pair of adjacent objects was 18° (e.g. standing at the dice, and pointing to the shoe and the lamp). The corridor lengths and angles were adjusted to fit this new layout while keeping the total path length similar to the possible maze (about 33.25m; corridor width remained 90cm). Likewise, the succession of left and right turns was kept constant across both mazes, ensuring a comparable memory load.

Starting at the book, the participant walked one lap through the impossible maze and ended up at the book again, but it was located at a different position in physical space, 6.23 m from the starting position (Figure 2d). Effectively, the participant was teleported across the gap in space to their initial point in the maze, which was achieved by instantaneously rotating the virtual environment 108° in the direction opposite to the walking direction. This transition was visually seamless, such that the view in the first corridor matched the view in the last corridor. This enabled the participant to continuously walk another lap. The local corridor-to-corridor metrics thus remained constant across each lap, but the positions of the objects in physical space, as given by idiothetic information, were globally inconsistent.

The impossible maze was slightly bigger than the walkable space in the tracking hall. To prevent collisions with the walls, we restricted the walkable area within the virtual world by placing red lines on the floor at a distance of 90cm from each acute corner in both the impossible and possible maze (Figure 2c). Participants were instructed to not walk the area behind those lines.

### Procedure

The experimental session started with a briefing about the procedure. Participants were informed that they first had to learn the layout of a virtual environment and the places within (green dot associated with an object) and that their spatial memory for that environment would be tested in a subsequent spatial task. They were told: “Your task is to memorize as accurately as possible the locations of these places in the environment. Later, in the test phase, in each trial you will be teleported to one place and asked to indicate the exact locations of the other places from your current position.”

### Learning phase

After donning the backpack and the HMD, the participant was disoriented by several random left and right turns (HMD display off) and led on a random path to the starting position. Then the virtual environment appeared, displaying the starting place, which was always the book. Additional verbal instructions were given at this point. In a pseudo-random fashion, half of the participants in each maze type group (nine each) began walking in a clockwise (cw) direction for three consecutive laps, before turning around and walking three laps in the opposite direction. The other half started learning the environment walking counterclockwise (ccw) first, before walking clockwise. They could walk at their own pace, but were restricted from looking backward. Only when they arrived at an alcove and stood right on top of a green dot they could take a full look around to visually explore the object, the corridor, and the connections to the two adjacent corridors. They were informed that the red lines on the floor should not be crossed, but the junctions could be used as visual cues to understand the angles between corridors. Participants could ask questions about the procedure throughout the experiment, excluding the nature and structure of the environment. During the first lap the participant was asked to name each object to ensure correct identification of the place labels.

After six laps, a learning check was carried out. The objects were removed and the participant was asked to walk two laps cw and two laps ccw while recalling the name of every other place they passed. Thus, all seven objects were queried in both directions. If an error was made, the participant repeated the learning procedure for two more laps, one cw one ccw, followed by another learning check. This process continued until the participant reached the learning criterion of 100% accuracy and moved on to the testing phase, or until six learning checks were unsuccessful, in which case the experiment was terminated.

### Testing phase

During a short pause of approximately five minutes the participant read written instructions for the testing phase. The test was conducted standing at a fixed position in the tracking hall. Pre-recorded audio instructions spoken by a male native English-speaker were used to guide the participant through the trials. Data collection was preceded by four randomly selected blocks of practice trials, which followed the same procedure.

Testing consisted of blocks of four ‘pointing’ trials, in either a clockwise or counterclockwise sequence. At the beginning of each block the participant was randomly teleported to one of the seven learned places, so they were standing on the green dot in the middle of the alcove. They could fully rotate their body around the spot but were instructed not to leave the green dot. The object and an audio recording (e.g., “You’re at the shoe”) provided information for self-localization and orientation. The view along the corridor was blocked by white fog (see Figure 3a), so only the object, the alcove, the back wall, and a few centimeters of the corridor on either side could be viewed. The participant was instructed to press a button on a handheld controller in response to the recorded question, “Do you know where you are?” to indicate that they were oriented. Thereupon the object in the alcove disappeared and a target object was named (e.g., “Face the lamp”). At the same time a vertical, black line appeared in the center of the field of view, which rotated with the participant’s head. The participant’s task was to ‘point’ in the direction of the target by turning to face the corresponding green dot, “as if the walls were made of glass”, thereby aligning their head and the black line with the remembered location of the target place. After they confirmed the facing direction with a button press, the next target was named, in order around the circle. Participants were instructed to face the target dot as accurately as possible with no time limit, and without feedback. After four consecutive targets, they were teleported to another learned place and the next block began.

**Figure 3.**
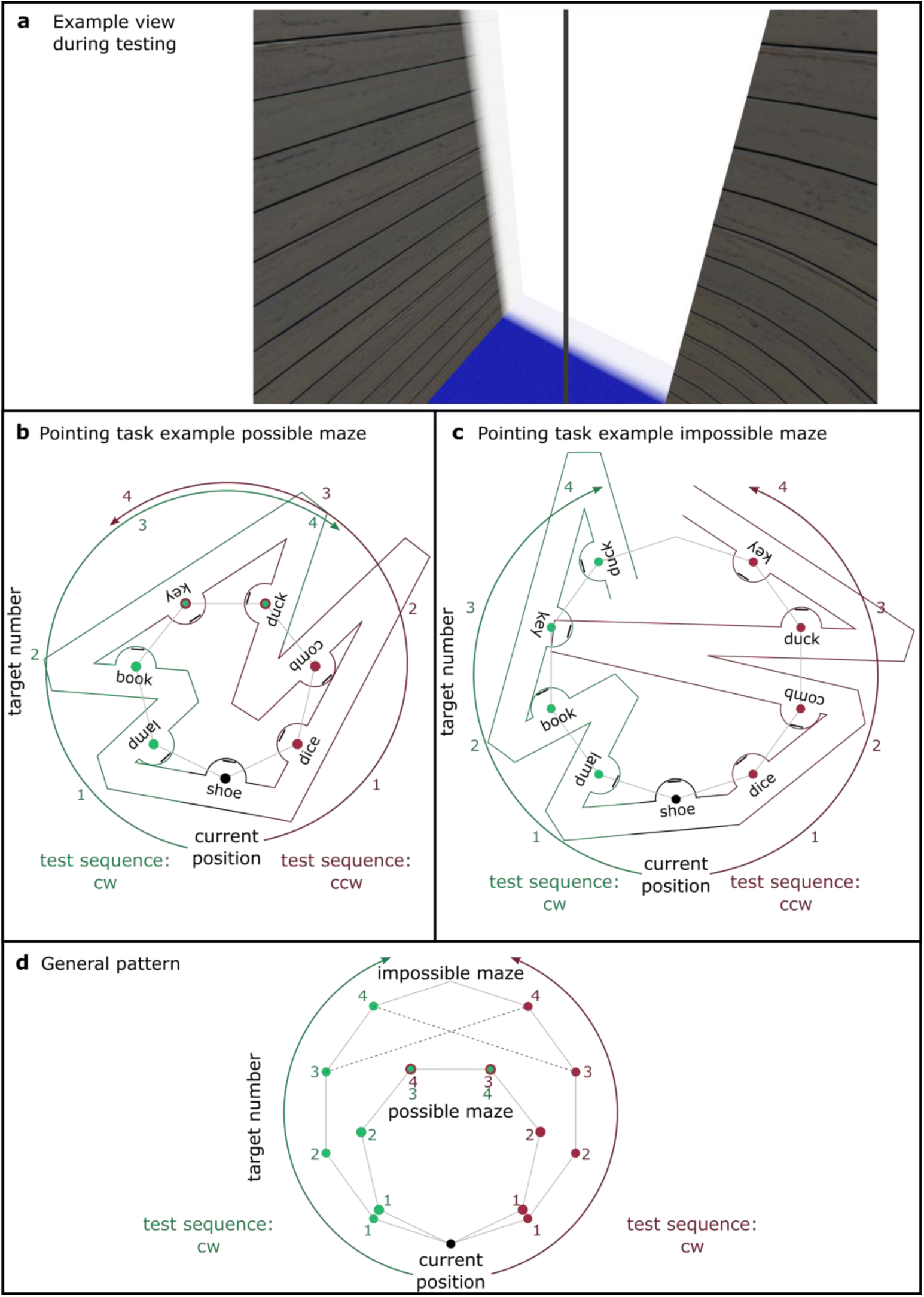
Testing phase. **(a)** Example of an egocentric view during testing when looking to the left while standing at the green dot of one’s current location. The view along the corridor was blocked to both sides of the alcove by a white fog. The black line at the center of a participant’s field of view (following every head movement) should be used as an aiming device to face the straight-line direction towards the target place. **(b) + (c)** Two example blocks of trials from the possible and the impossible maze. Standing at the shoe (black corridor) within one block of trials participants had to face four targets in a row, either along a clockwise (cw) test sequence (green corridors and arrow) with increasing target number (1-4), or along a counterclockwise (ccw) test sequence (red corridors and arrow). In the possible maze group (b) we expected that participants point out the same directions to the same target whether queried in cw or ccw test sequence. For the impossible maze (c) the local place-to-place metrics do not match up on a global scale. Depending on the order of test sequence (cw or ccw) the use of local metrics should lead to different estimated directions for the same targets. For example, the facing direction of key and duck (target number three and four) should yield average estimates shifted to the left for the cw test sequence compared to the ccw test sequence, just as if participants would point to two different locations. **(d)** General pattern of the target location based on local metrics relative to one’s current location for the possible (heptagon) and impossible (decagon) maze group. This pattern can be applied to any location participants were tested from because of the symmetry and even spread of the places across heptagon or decagon. Compared to the possible maze we expected an outwards bias for the impossible maze group when following local place-to-place metrics.

In each block of trials, participants had to face four targets in a predefined sequence in either the clockwise (cw) or counterclockwise (ccw) direction around the circle, starting with the nearest neighbor, followed by the adjacent neighbor and so on. An example is visualized in Figure 3b and 3c. Standing at the shoe, in the cw direction (green) participants first had to face the lamp, then the book, the key and finally the duck before being teleported to a new location. In another block, in the ccw direction (red), participants standing at the shoe first had to face the dice, then the comb, then duck and key, followed by teleportation. Participants were never tested around the full circle in consecutive order.

The experimental design was thus 2 *maze type* (possible vs. impossible) x 2 *test sequence* (cw vs. ccw) x 4 *target number* (1-4), with maze type a between-subject variable and the other two within-subject variables. From each of the seven learned places, there were two blocks of four ‘pointing’ trials (one cw and one ccw) presented in a random order, yielding 56 trial types. There were two repetitions, blocked separately, for a total of 112 trials per participant.

Participants were given short breaks of self-determined length after every 28 trials. Dependent measures included response latency and pointing error on each trial (angle between the facing direction and the target direction given by the local metrics vs. global embedding prediction).

### Debriefing

After the experiment we asked participants to draw a map of the environment and to fill out a questionnaire. In the questionnaire and in a subsequent debriefing we tried to assess whether participants in the impossible maze noticed the global inconsistencies by asking indirect (“Did you notice anything unusual in the environment?”) and direct questions (“Sometimes we visually teleported you from one object location to another. Did you notice?”). Sometimes participants reported the impression that the order or number of places or the length of the corridors changed during learning. As this was not the case, we attributed such comments to the difficulty of learning a complex environment rather than noticing a global inconsistency. Additionally, we marked down comments made by the participants during the learning phase, the testing phase or short interview in the debriefing phase that indicated detection of global inconsistency (e.g., “I should not be back at the book yet”; “Is this environment possible?”; “There are two places I could point to, where should I face?”, “I noticed something was off, but I tried to make sense of it.”). To validate their answers, we posed the same questions to the possible maze group. If either of these records suggested that a participant felt that the environment did not match up he/she was labeled as someone who “noticed” a mismatch. If not, he/she was denoted as “not noticed”.

### Data processing

As mentioned previously, four participants were removed from the impossible maze group and six from the possible maze group due to random pointing performance. For the possible maze group, pointing error was defined relative to the target’s Euclidean location. If mean absolute pointing error was not significantly better than chance (90°) we concluded that insufficient spatial learning had occurred and the participant was removed. For the impossible maze group, pointing error was calculated in two ways: relative to the location determined by the local place-to-place metrics (decagon), and relative to the Euclidean location for a global embedding with even spacing (heptagon). If the mean absolute pointing error was not significantly better than chance (90°) in either case, the participant was removed. From the remaining 36 participants we excluded 4.0% of the pointing error data and 4.9% of pointing latency data, as it deviated more than ±2 SD from a participant’s mean performance.

Since we were interested in biased pointing behavior predicted by local metrics, our analysis relied on the evaluation of the constant (signed) error (leftward and rightward biases) rather than the absolute error. In the impossible maze group 13 trials were identified where the participant exhibited pointing error close to 180° (facing away from the correct direction), so the sign of the error was ambiguous. These trials were excluded from analysis. Thanks to the symmetry of the underlying heptagon and decagon, the overall layout of targets is the same from each pointing position (Figure 3d). Thus, the pointing responses from each of the seven positions were combined for statistical analysis.

### Predictions

According to labeled graph theories, recall of a target location with respect to one’s current location should activate the sequence of stored place-to-place metrics between them. In a circular environment with seven places, there are two possible sequences to any given target, in the clockwise or the counterclockwise direction. Therefore, the manipulation of the cw or ccw test sequence was designed to bias the order in which local place-to-place information would be accessed. In particular, with test blocks of four consecutive targets, the last two targets overlapped in the cw and ccw direction (targets three and four). For example, in Figure 3b and 3c, when standing at the shoe, the last two targets in both directions were the key and the duck. In the possible maze, pointing responses to these targets should be approximately the same in both sequences (Figure 3b), but in the impossible maze, the local place-to-place metrics predict different responses when tested in the cw and ccw directions (Figure 3c). Specifically, pointing estimates in the impossible maze should show a strong outward bias, with errors to the left in the cw sequence, and to the right in the ccw sequence.

In contrast, according to metric map theories, local place-to-place measurements in the impossible maze should be embedded in a globally consistent Euclidean space during learning. We would thus expect the seven remembered target locations to approximate evenly spaced positions on a circle, similar to the symmetric heptagon. Of course, particular corridors might be susceptible to local deviations, which might differ across participants. But given that we combined data from the seven pointing positions, such local deviations should average out. Therefore, if participants learn a Euclidean map, they should point in the same direction to the same target, whether tested in the cw or ccw sequence, in both the impossible and possible mazes.

### Transparency and openness

We report how we determined our sample size, all data exclusions, all manipulations, and all measures in the study. All data are available as supplementary material. Materials and analysis code for this study are not available. This study was not preregistered.

## Results

On average participants in the possible maze walked 8 laps (*SD* = 3.20) during learning and performed 2.00 (SD = 1.60) learning checks to reach the learning criterion of 100% correctly recalled places. They spent 32.30 min (*SD* = 20.87) learning the environment, and a total of 43.89 min (SD = 22.81) in the environment including the checks. Correspondingly, in the impossible maze participants walked 6.63 laps (*SD* = 1.34), did 1.32 (SD = 0.67) learning checks, spent 24.89 min (*SD* = 9.39) learning the environment, and a total time of 32.63 min (SD = 12.61) including checks. Neither the learning time nor the total time was significantly different between groups, *t*s < 1.89, *p*s > .070.

### Clockwise vs counter-clockwise test sequence

If pointing responses are based on a Euclidean map, participants should point in the same direction to the same target when tested in the cw and the ccw direction, in both mazes. In contrast, if a graph-like representation underlies pointing estimates, leftward biases are expected for the cw sequence, and rightward biases for the ccw sequence, in the impossible maze. Therefore, in our first analysis we examined whether participants’ pointing directions to the two overlapping targets (targets three and four) deviated from a symmetric heptagon. The predictions are summarized in Figure 4a.

**Figure 4.**
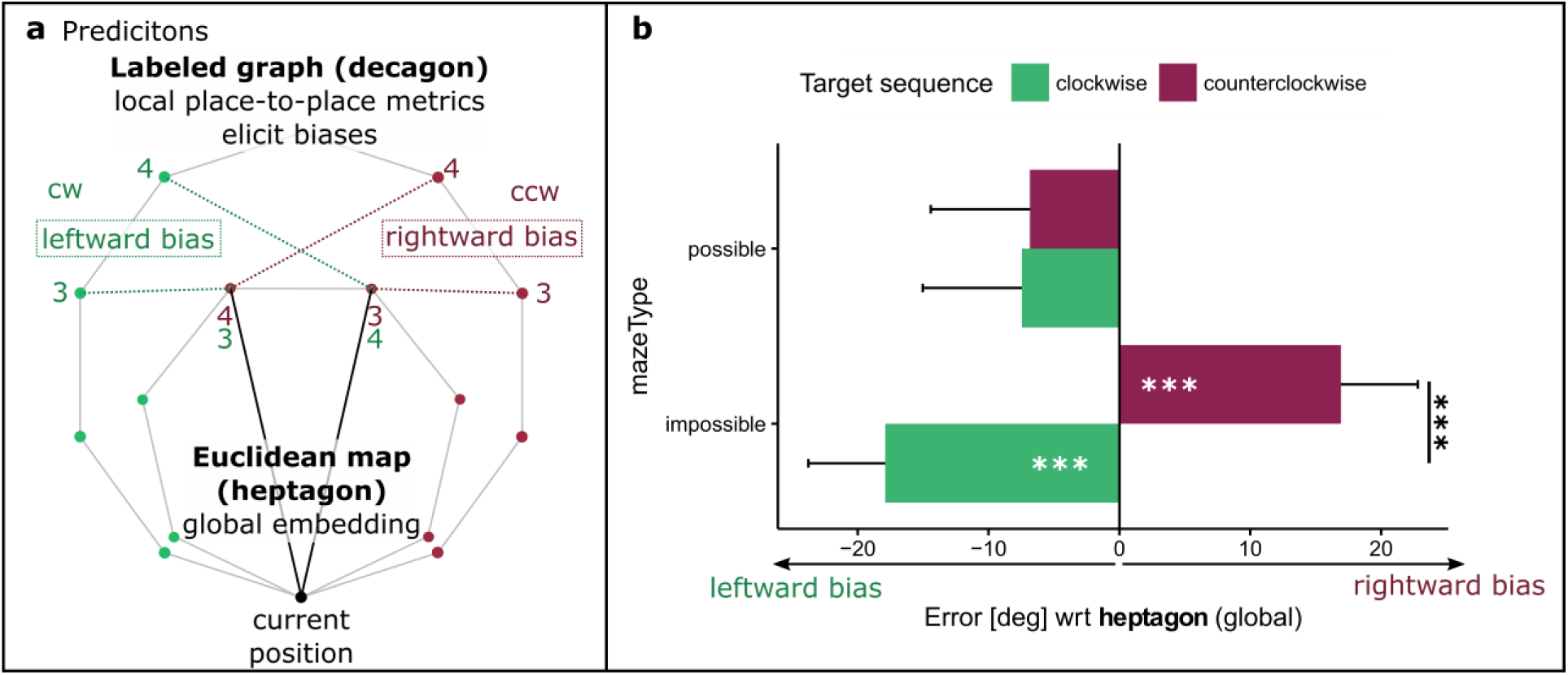
**(a)** Predicted pointing patterns when facing the same targets (target number three and four) in the cw or ccw sequence. If participants learn a global embedded Euclidean map, the impossible maze group should point in the same direction independent of test sequence (cw or ccw), similar to the possible maze group (heptagon). If they rely on a graph-like representation of local place-to-place metrics, the impossible group should exhibit leftward and rightward biases in the cw and ccw test sequences, respectively. **(b)** Mean pointing error for targets three and four relative to a symmetric heptagon, in the possible and impossible mazes. Whereas the possible maze group pointed in the same direction irrespective of the test sequence, the impossible maze group exhibited clear leftward and rightward biases for the same targets, pointing in significantly different directions when queried in a cw or ccw sequence. Error bars indicate within-subject standard errors. * p < .050; *** p < .010

We first computed the constant (signed) pointing error with respect to the symmetrical heptagon predicted by a global embedding, where a value around 0 indicates pointing towards the heptagon locations (i.e., global embedding), a positive value indicates a rightward error and a negative value a leftward error. Results for targets three and four are depicted in Figure 4b, where errors in the possible maze appear on the top and the impossible maze on the bottom. We performed a mixed two-way ANOVA with the factors *maze type* (possible vs. impossible) and *test sequence* (cw vs. ccw) on the data from targets three and four. There was no overall effect of *maze*, *F*(1, 34) = 1.42, *p* = .241, *η_p_^2^* = .04, but we observed a main effect of *test sequence, F*(1, 34) = 6.84, *p* = .013, *η_p_^2^* = .17, as well as a significant interaction, *F*(1, 34) = 6.37, *p* = .016, *η_p_^2^* = .16. Post-hoc t-tests found no difference between the cw and ccw sequence in the possible maze, *t*(34) = -0.06, *p* = .949, but a significant difference in the impossible maze, *t*(34) = -3.63, *p* < .001. Specifically, the cw test sequence yielded a leftward bias that was significantly different from zero, *t*(17) = -2.93, *p* = .009, while the ccw sequence yielded a significant rightward bias, *t*(17) = 4.03, *p* < .001. Thus, the impossible maze group did not point in the same direction in cw and ccw trials, despite being queried to point to the same targets. This pattern was identical for target three and target four when examined separately, revealing leftward and rightward biases in the cw and ccw directions, respectively, *p*s < .010.

Based on their responses in the debriefing, eight participants in the impossible maze group were classified as having “noticed” an inconsistency in the environment, and the remaining ten participants as having “not noticed” anything. An ANOVA for the impossible maze group with the between subject factor *awareness* (noticed vs. not noticed) and the within-subject factor *test sequence* (cw vs. ccw) revealed neither a main effect of nor an interaction with *awareness*, *p*’s > .103. Only the main effect of *test sequence* was significant, *F*(1, 16) = 16.76, *p* = .001, *η_p_^2^* = .51, confirmed by post-hoc tests of participants who “noticed”, *t*(16) = -3.11, *p* = .007, and those who did “not notice”, *t*(16) = -2.72, *p* = .015. Thus, participants in the impossible maze were significantly biased by the cw or ccw test sequence whether they had noticed inconsistencies in the environment or not. Interestingly, four participants in the possible maze group also reported noticing some inconsistency (e.g., “I felt I was moved to a different position each lap.”, “I couldn’t make a circle in my mind.”) despite learning a Euclidean environment.

### Pattern of pointing errors

Second, we analyzed the pointing directions in the impossible and possible maze, to determine whether the outward biased error pattern was predicted by the local place-to-place metrics experienced during learning. For this analysis, we collapsed the data in the cw and ccw test sequences by multiplying the constant errors on cw trials by -1, so a positive error represents an outward bias and a negative error represents an inward bias relative to the heptagon locations^1^. Values approximating zero represent pointing to the heptagon layout, corresponding to the prediction of the global embedding hypothesis in the impossible maze, and both hypotheses in the possible maze. Taking the heptagon locations as a reference, we can also make precise predictions for the labeled graph hypothesis in the impossible maze, based on the local place-to-place metrics of the decagon (Figure 5a). This yields an expected outward bias of ca. 7.71° for target one, and an additional bias of ca. 7.71° for each successive target up to ca. 30.86° for target four. The graph hypothesis thus predicts a positive linear relationship between target number and outward bias in the impossible maze.

**Figure 5.**
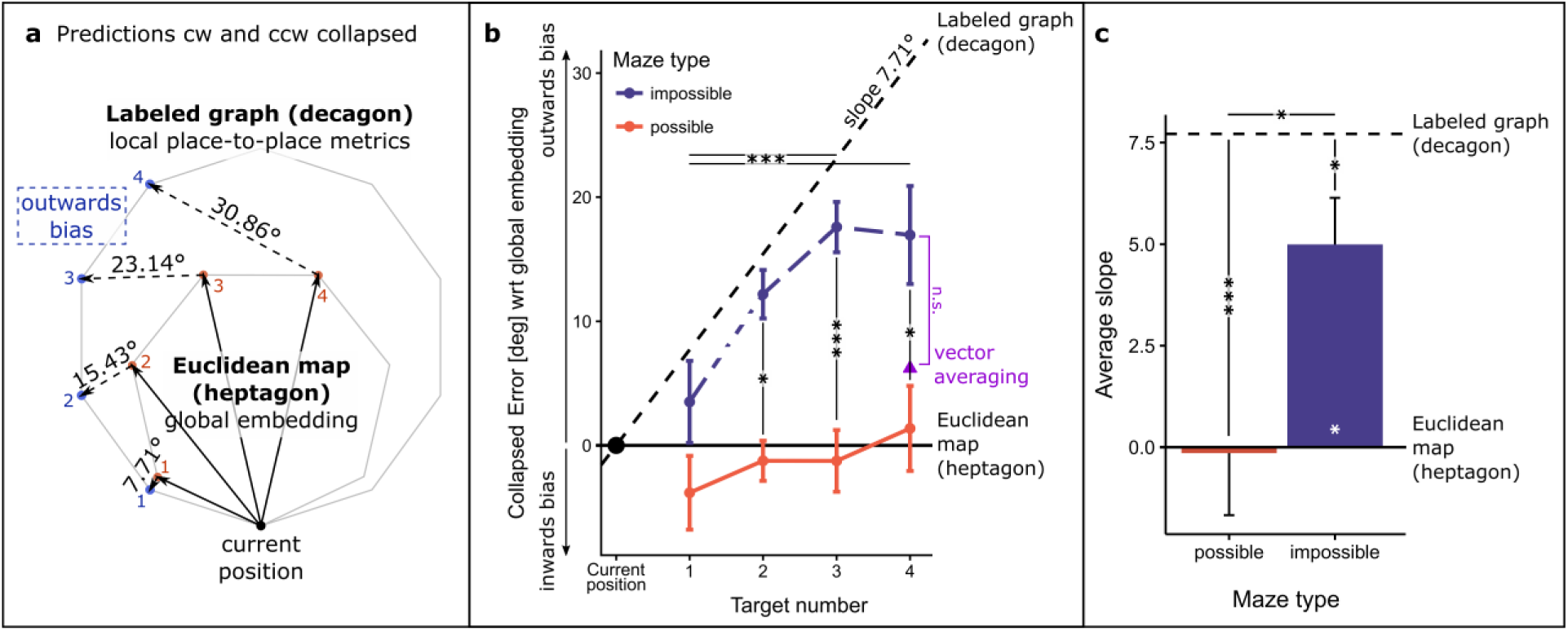
Examination of outward biases. **(a)** Predicted outward biases relative to a global embedding when instead following the local place-to-place metrics. Matching up the heptagon and decagon underlying the target layouts for possible and impossible maze a linear increase of the outward bias by ca. 7.71° per additional target number in a block can be expected in the impossible maze group if pointing is based on a local graph representation. **(b)** Collapsed error (positive values representing an outward bias) across target numbers and the predicted patterns for either the Euclidean map (estimates approaching a heptagon) in black solid line (zero line) or the local graph representation (estimates following vertices of decagon) in black dashed line with a slope of 7.71°. **(c)** Average slopes extracted from linear regressions of individual performances across target number. Average slope of the impossible maze group leans towards the predicted slope for a graph representation but remains significantly different from it. *t*(71.16) = 2.25, *p* = .027, three, *t*(71.16) = 3.16, *p* = .002, and four, *t*(71.16) = 2.61, *p* = .011, although not target one, *t*(71.16) = 1.23, *p* = .223, given the small expected difference of only 7.71°. Furthermore, a main effect of *target number* was found, *F*(1.89, 64.15) = 4.38, *p* = .018, *η_p_^2^* = .11, but the interaction was marginal, *F*(1.89, 64.15) = 1.48, *p* = .237, *η_p_^2^* = .04 (the Greenhouse-Geisser adjustment was applied to within-subject comparisons). Post-hoc Tukey tests comparing error patterns across target numbers revealed significant differences between targets only in the impossible maze group: outward bias increased significantly from targets one to three, *t*(102) = -3.53, *p* = .004, and one to four, *t*(102) = -3.37, *p* = .006 (remaining *t*s ≤ 2.17, *p*s ≥ .136). No target differences were observed in the possible maze group, *t*s ≤ 1.30, *p*s ≥ .565.

The results appear in Figure 5b, which represents outward bias as a function of target number, where the global embedding hypothesis (heptagon) predicts a line with a slope of zero (solid line), and the labeled graph hypothesis (decagon) predicts a line with a slope of 7.71° and an intercept of zero (dashed line) in the impossible maze. The y-intercept of zero represents the current pointing position. A 2-way mixed ANOVA on outward bias with the factors *maze* and *target number* confirmed that the error patterns of the possible and impossible maze groups differ, supporting our first analysis. There was a main effect of *maze F*(1, 34) = 8.07, *p* = .008, *η ^2^* = .19, with post-hoc t-tests showing significant differences between groups for targets two, We again checked for a difference between participants who “noticed” and those who did “not notice” a discrepancy in the impossible maze. A 2-way ANOVA (*awareness* x *target number*) on outward bias found neither a main effect of *awareness*, *F*(1, 16) = 3.38, *p* = .085, *η_p_^2^* = .17, nor an interaction, *F*(1.81, 29.01) = 1.45, *p* = .250, η_p_^2^ = .08, only a main effect of *target number*, *F*(1.81, 29.01) = 5.04, *p* = .016, *η_p_^2^* = .24. Noticing a discrepancy thus does not influence the pattern of errors.

To examine whether the increase in error with target number is in accordance with the slope predicted by local place-to-place metrics, we performed a linear regression on each participant’s data. The y-intercept, corresponding to participant’s pointing position, was forced to be zero. The mean slope for the possible and impossible maze groups appears in Figure 5c, showing a significant difference between them, *t*(31) = 2.69, *p* = .011, *d* = .90. The average slope of 5° per consecutive target in the impossible maze group is significantly higher than the slope of 0° expected under the global embedding hypothesis (corresponding to the heptagon), *t*(17) = 4.37, *p* ≤ .001. Yet it is also smaller than the slope of 7.71° expected under the labeled graph hypothesis (corresponding to the decagon), *t*(17) = -2.38, *p* = .029. The average slope for the possible maze group of -0.15° is not significantly different from the expected slope of 0° for this group, *t*(17) = -0.10, *p* = .925, reflecting a good fit to the heptagon prediction; this slope is also significantly smaller than 7.71°, *t*(17) = -5.14, *p* ≤ .001.

To compare the two hypotheses in the impossible maze group, we adopted a Bayesian model selection approach, specifically comparing two different linear regression models of the relationship between target number and outward bias: (1) the labeled graph model with a slope of 7.71° and (2) the Euclidean map model with a slope of 0°. The regression was performed on the mean outward pointing error for all participants (the data underlying Figure 5b), forcing the intercept to zero. To calculate the likelihood of each model, sigma was based on the standard deviation of error in the possible maze control group (*SD* = 19). The Bayesian Information Criterion for the labeled graph model (BIC_LG_ ) was 639.1, and that for the Euclidean map model (BIC_EM_) was 647.0, yielding a difference of ΔBIC = BIC_EM_ – BIC_LG_ = 7.9. We further calculated the Bayes Factor to compare the two models, yielding BF_GM_ = 52.9. The size of both the ΔBIC and the BF_GM_ indicate very strong support for the labeled graph model over the Euclidean map model (Jeffreys, 1961).

### What is happening at target four?

As can be seen in Figure 5b the impossible maze data closely follow the prediction of the local graph model for targets one to three, *t*s < 1.53, *p*s > .144, but flatten out at target four, which differs significantly from the graph prediction, *t*(17) = 2.58, *p* = .019. However, it also clearly differs from the global embedding prediction, *t*(17) = 3.15, *p* = .006. Target four is unique because, although it is the fourth target in the biased sequence, it is only the third target in the opposite, non-biased sequence. For example, following a cw target sequence, the fourth target corresponds to the third target in the opposite ccw sequence. In this case, the “mental route” through a labeled graph from the pointing position to the target is shorter in the ccw direction than the cw direction. Consequently, if a labeled graph structure underlies spatial localization, participants might sometimes follow the shorter ccw sequence to target four, with fewer nodes and a smaller path length, instead of continuing to activate nodes in the cw sequence. The logic is the same for the reverse example of following a ccw target sequence.

To investigate this possibility, we performed a number of exploratory analyses on the data for target four in the impossible maze, as explicated in the Supplementary Materials. Briefly, we failed to find a spike in the mean within-subject SD that would suggest a participant followed the cw sequence on some trials and the ccw sequence on others when estimating target four. Similarly, we found no evidence of increased pointing latency for target four that would suggest a change in estimation process. Also, there was no evidence of a bimodal distribution in pointing error between-subject, which would suggest that some participants followed the biased and others the unbiased sequence for target four. There was an increase in the between-subject SD, but it was due to a few participants who showed very large outward or inward biases. Finally, it is possible that the pointing direction to target four was estimated in both the cw and ccw order and then averaged before making a response, although this should also have increased the latency to target four. Alternatively, responses may have been biased toward the center of the alcove when there was competition between the cw and ccw directions. If the former occurred on every trial, it predicts an outward bias of 6.21° (triangle in Figure 5b), which was not statistically different from the observed bias of 17.4°, *t*(17) = 1.27, *p* = .223. Pointing always to the center of the alcove predicts an even larger outward bias of 12.86°.

### Are the possible and impossible mazes comparable?

To compare the process of target localization in the possible and the impossible mazes, we examined the within-subject variability in pointing direction, as well as the pointing latency, for all four targets. First, we performed a 2-way mixed ANOVA on the standard deviation of individual pointing bias. There was a main effect of *target number, F*(1.86, 63.29) = 7.68, *p* = .001, η_p_^2^ = .18, but no main effect of *maze type*, *F*(1, 34) = 0.02, *p* = .877, *η_p_ ^2^* < .02, nor an interaction, *F*(1.86, 63.29) = 0.47, *p* = .616, *η_p_^2^* = .01. For both maze groups variability in performance increased with target number. Post-hoc t-tests yielded significant differences between targets one and three, *t*(102) = -3.17, *p* = .011, one and four, *t*(102) = -4.40, *p* ≤ .001, and two and four, *t*(102) = -3.15, *p* = .011, remaining *t*s ≤ 1.94, *p*s ≥ .222. The increase in SD can be explained by the accumulation of error due to the successive activation of consecutive nodes in the target sequence, as predicted by approximate place-to-place metrics in a labeled graph. Note that variable error increased linearly with the path length through the graph, rather than the straight-line distance to the target, which was a decelerating increase for targets one to three and equal for targets three and four.

Second, we ran a 2-way ANOVA on pointing latency. Only a main effect of *target number* was found, *F*(1.43, 48.59) = 23.68, *p* < .001, *η_p_ ^2^* = .41, but no main effect of or interaction with *maze type*, *F*s ≤ 0.66, *p*s ≥ .422. In both mazes, pointing to target one took significantly longer than pointing to the succeeding targets, *t*s ≥ 6.11, *p*s ≤ .001, and differences between targets two to four were not significant, *t*s ≤ 1.09, *p*s ≥ .698. This pattern is consistent with the successive activation of nodes in a labeled graph during a block of four consecutive targets. Thus, pointing latency was comparable in both maze groups.

### General Discussion

The present study set out to test two hypotheses about the memory structure that underlies survey knowledge of a complex environment. The metric map hypothesis proposes that locations in space are assigned to coordinates in a Euclidean coordinate system, yielding globally consistent spatial knowledge. In comparison, the labeled graph hypothesis suggest that we store local information only, such as the approximate distance and direction between adjacent places. These local place-to-place metrics depend on experience and incorporate error, and thus may be globally inconsistent. Consequently, variation in how the individual places are recalled during navigation can result in different estimates of the distance and direction to the same target.

We contrasted these hypotheses by asking participants to learn a seven-corridor zig-zag loop, which was either a possible Euclidean environment or an impossible non-Euclidean environment in which the local place-to-place metrics were globally inconsistent. The possible maze group pointed consistently to the same targets in a heptagonal pattern, irrespective of the target sequence during testing. In contrast, the impossible maze group did not point in the same direction to the same target. Instead, participants exhibited significant leftward and rightward errors, depending on whether they were tested in a cw or a ccw order around the loop. These pointing errors reflected an outward bias expected when relying on the local place-to-place metrics of a graph. The pattern of this bias is better explained by the labeled graph hypothesis than the global metric map hypothesis, which assumes the place-to-place measurements are embedded in a geometrically consistent Euclidean space, regularizing the local biases.

### Violation of the metric postulates

The results conflict with the metric postulate of positivity, which states that the distance from any point to itself must be zero. Consequently, each unique place can only be assigned to a single location in physical space, and no other place can be assigned the same coordinates (e.g., Beals et al., 1968; McNamara & Diwadkar, 1997). Our first analysis showed that when the impossible maze group was asked to point to the same target, however, it was remembered as occupying two widely separated locations in physical space. Specifically, when asked to point to target three (or target four), they pointed to locations that were, on average, 34.8° apart. This result confirms Warren, et al’s. (2017) finding of shortcuts that indicate large violations of the positivity postulate in spatial knowledge.

Previous reports of violations of the symmetry postulate (e.g., Burroughs & Sadalla, 1979; McNamara & Diwadkar, 1997; Sadalla et al., 1980) have been countered by two bias models in which asymmetries are not ascribed to the spatial representation but to biases in the estimation process. The category-adjustment model of spatial coding (Huttenlocher et al., 1991; Newcombe et al., 1999) assumes representing space on a fine-grained (conceivably Euclidean) level and a coarse-grained categorical level. Biases are produced during retrieval by adjustments made to the fine-grained place information based on its distance to a category prototype. For example, a typical category prototype is the middle of a quadrant of a circle which is drawn on a piece of paper and formed by vertical and horizontal visual axes. Location estimates are biased toward the center of the category, resulting in asymmetric biases. Alternatively, the contextual-scaling model (McNamara & Diwadkar, 1997) assumes that different places evoke different contexts in working memory when being referenced, depending on their salience (e.g., familiarity, functional importance). Thus, a different context is activated when standing at place A recalling the location of B compared to standing at place B recalling A. This scales the retrieval process accordingly and leads to asymmetries. Both models allow for the existence of a Euclidean mental map and render distance asymmetries less a violation of metric postulates than originally thought.

Can these models explain the present results as well? We believe they cannot. In the models the asymmetry arises from interchanging either the reference location or the category prototype. Our study, however, compares blocks of trials that have the same reference object (i.e., current position of participant) and the same target object, and only varies target order (cw vs. ccw). One and the same target object relates to the same categorical bias according to the category-adjustment model and one reference object relates to the same location context according to the contextual scaling model.

A second argument can be made against the explanatory power of the two models with regards to our study. First, given that we combine all blocks of trials for analysis (e.g., merging blocks in which the participant points from the book to the dice and from the dice to the book), all potential context and category effects should average out. Yet we still observe diverging pointing directions to the same target. The divergence is also not an artifact of this averaging process, for the leftward and rightward bias in cw and ccw test sequences was observed from each pointing position. For four out of the seven reference objects (i.e., pointing positions) the difference in pointing direction was statistically significant, and the remaining three went in the predicted direction (additional analysis not reported in the results). Thus, despite activation of the same context or category prototype (i.e. constant reference-target pairings), the direction estimates still differed.

One might argue that the context-scaling model could be adjusted by adding the assumption that previous estimates of targets one and two are considered for subsequent estimates as well. Each estimated target possesses its own saliency (note that differences in saliencies between places should be averaged out as all blocks of trials are merged). Each previous target might bias the subsequent estimates towards their location. We do not think this is a likely explanation for our results, as in this case we should see similar outward biases in the possible maze group as well.

We conclude that current bias models are insufficient to explain the present findings. Our results agree well with previously reported asymmetries in direction estimates performed forward along a learned route compared to route backward, which are independent of the reference and target objects (e.g., Meilinger, Henson, Rebane, Bülthoff, & Mallot, 2018; Meilinger, Strickrodt, et al., 2018). These route direction effects are also difficult to explain with bias models. Correspondingly, the present results suggest that we successfully probed place-to-place metrics in a labeled graph by manipulating the target test sequence.

### The labeled graph hypothesis

In our second analysis, we ascertained that the outward bias observed in the impossible maze was closely predicted by the local place-to-place metrics for the first three targets. This finding supports the view that spatial knowledge reflects the local information that is experienced during learning, rather than embedding such measurements in a globally consistent Euclidean map. Over all four targets, the present results strongly support the labeled graph hypothesis over the metric map hypothesis.

Our results also lend support to the interpretation of previous studies using impossible mazes as evidence against Euclidean mental maps. In particular, the wormhole experiments of Warren and colleagues (Ericson & Warren, 2020; Warren et al., 2017) demonstrated large inconsistencies in spatial knowledge that were contradictory to a metric map, but consistent with a labeled graph. The present study supports that interpretation, and goes further by showing that the magnitude of directional biases can be predicted by the local place-to-place metrics. Taken together, these findings indicate that a labeled graph structure incorporating local metric information best explains current and previous data.

Previously, Kluss and colleagues (2015) reported that that the local angles in an impossible triangle environment were reproduced as they were experienced and not embedded in a globally consistent Euclidean structure, in which the inner angles must sum to 180°. However, this report was not substantiated statistically, for the sum of inner angles was not significantly different from 180°. Indeed, we faced a similar problem in a preliminary study for the present experiment, based on possible and impossible versions of a hexagonal six-corridor maze. The corridors had a much simpler and more regular layout than in the present study, and the impossible version was only a slightly distorted hexagon “opened up” to fit six vertices of a heptagon. Consequently, the predicted difference in outward bias between the two mazes was too small to be measured reliably. Despite notable biases in the error patterns, the differences were not statistically reliable, given the variability in pointing responses. Hence, future research should be careful to design environments that allow the detection of differences with typically noisy survey tasks.

### Sequential activation in a labeled graph

Importantly, previous studies have shown that the number of corridors between the pointing location and the target is correlated with the latency of the estimate (e.g., Meilinger et al., 2016; Meilinger, Strickrodt, et al., 2018; Strickrodt et al., 2018). Such corridor distance effects cannot be explained by the straight-line Euclidean distance between locations but are consistent with successive processing of places along a mental route through a graph linking the pointing location to the target location. Each additional node (e.g., a corridor) on the route is activated in sequence, thus incrementally increasing the latency to make the final estimate.

The present study both exploited and provided evidence for this process of piecewise activation. First, we assumed that successive activation during a block of four targets would prime the direction taken through a mental graph (cw or ccw) when making a pointing estimate. Indeed, the predefined sequence of targets yielded outward biases that reflected the local place-to-place metrics in the impossible maze.

Second, previous studies have shown that when pointing to a series of neighboring targets subsequent pointings seem to be based on previous pointings (Meilinger, Strickrodt, et al., 2018). More precisely, when pointing to a sequence of adjacent neighbors, only the difference between the new target and the previous target is calculated, rather than beginning the recall process at the pointing position each time, discarding the previously retrieved information. This process is reflected in our results as well, for both the possible and the impossible maze. Although the path length from pointing position to target location increased with target number, the pointing latency remained stable after the first estimate. This implies that for every new target, only one additional node was activated to estimate the target’s direction. Hence, subsequent estimates are not independent. Only after being teleported to a new pointing position at the beginning of a block of trials did this incremental estimation process begin again from scratch. Note that the longest response latencies were found for the first target within a block. The first trial has largest uncertainty about whether the cw or the ccw neighbor will be named as the target. As soon as this first estimate is completed, the next three targets can be predicted uniquely, and the direction of the first target serves as a reference for estimating the direction of the next target more rapidly.

A third argument for sequential activation of nodes in a graph is based on the within-subject standard deviation of pointing error. Individuals’ standard deviations increased linearly with target number in both in the possible and the impossible maze. Such a pattern can be explained by error accumulation during recall. Given some noise in local place-to-place information, the successive activation of nodes adds random error into the pointing estimate. With a Euclidean map, on the other hand, pointing estimates between the coordinates of the current position and the target are computed from scratch on each trial, without relying on the preceding estimates. Each response should thus be independent, and noise should not accumulate during a block. It is possible that pointing error might increase with the straight-line distance to the target, but the within-subject SD increases linearly with target number (path length through the graph), rather than target distance.

Taken together, the observed outward bias, the latency pattern within a block of pointing trials, and the linear increase of variability with the graph distance to the target are consistent with sequential activation of graph nodes during the pointing.

### The metric embedding hypothesis

A process of global metric embedding might occur at the time of encoding or during long-term memory consolidation. As an example of the former approach, Wang (2016) proposed that the path integration system continuously updates a vector pointing to each of the objects visited during learning. Applied to the current experiment, consider the example in the impossible maze illustrated in Figure 2b. Starting at book_1_, imagine walking cw to the lamp, just before returning to the book, now located at book_2_. Standing at the lamp, the updated vector to the starting point (book_1_) would be off by about 54° relative to book_2_. The same holds for the other objects. For every loop around the environment the updated vector to the previous location of an object will be off by about 54° when revisiting it in the next round. These vectors might then be adjusted to minimize the error over one lap, yielding an object layout that approximates a symmetric heptagon. As an example of the latter approach, Mallot and Basten (2009) describe how a global embedding could be achieved based on local information stored in long-term memory. The place-to-place metrics are compared by triangulation and adjusted to remove geometric inconsistencies.

The integration of local measurements in the impossible maze to construct a symmetric heptagon might appear to be a rather strong claim for the global embedding hypothesis. Yet we maintain that such procedures are theoretically possible in our seven-corridor environment. Each object was repeatedly visited in a loop at least six times during learning, providing sufficient exposure to acquire local place-to-place metrics. Participants were instructed to learn the environmental layout and told they would be tested on the locations of all places. The circular nature of the object array, with equal inter-object distances, placed strong constraints on a metric embedding, such that object locations must be represented on a circle with roughly equal spacing. Although local distortions presumably occurred across corridors and participants, they would tend to average out across multiple pointings from all places, so the resulting map of object locations should, on average, approximate a symmetric heptagon. We thus consider the symmetric heptagon to be a fair prediction of the global embedding hypothesis in both the impossible and possible mazes.

Nevertheless, our results show that pointing was driven by the local metrics, rather than a consistent global embedding. It is possible that vector updating by path integration might be ineffective in our multi-corridor environment, considering that previous studies supporting continuous updating were conducted in open environments (e.g., Jayakumar et al., 2019; Tcheang et al., 2011). Leaving a local environment was also found to impair the spatial updating of objects left behind (Wang & Brockmole, 2003). It is also possible that our environment and the required calculations were too complex for a mental triangulation process. However, such conditions are common enough in the real world, and the claim that they interfered with the construction of a geometrically consistent map would cast doubt on the robustness of a global embedding process.

### Awareness of maze inconsistencies

In contrast to previous virtual maze experiments, we seem to have created a geometric inconsistency that was large enough to be noticed by nearly half the participants. In the wormhole study by Warren and colleagues (2017), only one of 31 participants noticed a discrepancy, even though the environment rotated 90° and teleported them 6-10 m. Suma and colleagues (2012) found that roughly 14% of participants noticed that symmetric rooms connected by a straight corridor overlapped in an impossible manner, depending on room size and the amount of overlap. In our experiment, the impossible maze underwent a rotation of 108° in each lap, which accumulated over three succeeding laps during learning. In addition, the junctions in the impossible maze were visually elongated to clearly specify the local angle between corridors, but this could have heightened the impression that some corridors overlapped; in particular, the junction between duck and comb was very long and would have crossed over the opposite corridor in a regular heptagon (Figure 3c). Note, however, that while there were eight *noticers* in the impossible maze group, there were also four *noticers* in the possible maze group, even though the maze was actually Euclidean. Thus, the fact that the experiment took place in a complex, unfamiliar virtual environment may be partially responsible for this impression.

Nevertheless, this presented an opportunity to test whether awareness of a geometric inconsistency promoted a global embedding to achieve consistency. Interestingly, we found no evidence that this was the case, for there were no significant differences between *noticers* and *non-noticers* in the impossible maze. *Noticers* did not adjust their pointing estimates to be globally consistent, but continued to rely on local place-to-place metrics. Thus, even awareness of a discrepancy did not facilitate global embedding, and the pointing responses of both groups were consistent with a labeled graph.

Taken together, the present results imply that the structure of survey knowledge can be characterized as a labeled graph. Pointing estimates involve the sequential activation of nodes corresponding to places along a mental route through the graph from one’s current location to the target. This activation persists long enough to support subsequent estimates. Importantly, the metrics stored in this graph appear to be purely local and are not integrated to form a global, geometrically consistent map.

### Consideration of possible objections

A number of objections might be raised to the present study. First of all, having participants learn a non-Euclidean environment violated the geometric constraints of the real world and may have altered the normal spatial learning process. Perhaps participants learned a labeled graph of the impossible maze, but formed a Euclidean map of the possible maze – and the real world. We don’t believe that this is likely, however. If fundamentally different processes take place in Euclidean and non-Euclidean environments, resulting in qualitatively different knowledge structures, this should be reflected in the performance of the possible and impossible maze groups. First, during learning, the average number of laps and total time to criterion were comparable in the possible and impossible mazes. Second, in the test phase, there were no significant differences in within-subject SD or pointing latency between groups, indicating comparable precision in spatial locations. In particular, the within-subject SD increased with target number in the possible as well as the impossible maze, strongly implying a similar process of sequential place-to-place activation and error accumulation in both mazes. Thus, our results do not support the claim that spatial learning or spatial knowledge differ in Euclidean and non-Euclidean environments.

Another objection might be that participants were exposed to the maze for only half an hour in one session. This may have been sufficient to learn local place-to-place relations, but perhaps with more experience participants might eventually acquire a consistent Euclidean map. We cannot exclude this possibility, of course, but it is hard to test in practice because the argument that additional experience might alter the representation always holds, whether there are one or ten learning sessions. Nevertheless, we point out that the absolute error as an indicator of accuracy was within the range of errors observed after extended experience in real environments in previous studies (e.g., Ishikawa & Montello, 2006; Moeser, 1988; Schinazi, et al., 2013). An average of 44.81° (SD=16.81) absolute error was observed in the impossible maze and 45.32° (SD=17.74) in the possible maze.

Some previous studies have reported alignment effects which suggest that spatial information about multiple places and corridors can be integrated into a coherent format (e.g., Strickrodt et al., 2018; Tlauka et al., 2011; Wilson & Wildbur, 2004; Wilson et al., 2007). Specifically, when the participant is aligned with the global main orientation of the environment (e.g., the first perspective experienced), a survey task such as pointing is facilitated, suggesting that local information may be embedded in a global map. Yet effects of sequential place-to-place activation still remain (see also Meilinger et al., 2013; Strickrodt, et al., 2018), implying that the participant can’t simply read out map coordinates. Strickrodt and colleagues (2018) offered an alternative explanation, namely, that a general reference direction or vector may be associated with each node of the graph representation, specifying a common direction across local contexts. Despite these open questions, our results emphasize two points. First, a local labeled graph seems to characterize the basic structure of survey knowledge and second, a global Euclidean map is not necessary to perform survey tasks like straight-line pointing or taking short-cuts.

## Conclusion

The present results indicate that similar processes underlie survey estimates in both the possible and the impossible maze. At the same time, pointing directions in the impossible maze strongly violate the positivity postulate: when asked to point to the same target, the estimated direction depended on the preceding cw or ccw target sequence, and differed by 34.8° on average. Clearly, the impossible maze was not embedded into a geometrically consistent framework such as a Euclidean map. Instead, the observed outward bias in pointing was predicted by the local place-to-place metrics, corresponding to a labeled graph of the environment. Pointing responses are consistent with sequential activation of local nodes, analogous to a “mental route” through the graph. In sum, our results indicate that knowledge of navigable space consists of local piece-wise, potentially inconsistent information best characterized as a labeled graph (Chrastil & Warren, 2014; Meilinger, 2008; Warren et al., 2017), rather than a mental coordinate system that integrates the metric relations between places in a geometrically consistent Euclidean map (e.g., Byrne et al., 2007; Gallistel, 1990; O’Keefe & Nadel, 1978).

## Author notes

We would like to thank Joachim Tesch and Adam Hersko-Ronatas for help with the virtual environment setup and Gregory Dachner and Lauren Franklin for the audio recordings. All data are available as supplementary material. Materials and analysis code for this study are not available. This study was not preregistered.

## Supplementary Material

### What is happening at relative corridor distance four?

#### 1. Is there a higher variability in the pointing behavior within subject?

Do individual participants vary in the way they recall the fourth target? For example, sometimes they might be taking the predefined, biased “mental route”, sometimes they might switch to recalling the fourth target by successively activating local entities along the non-biased direction. If this is the case participants should show higher variability in their error pattern for corridor distance four and the time needed for facing should increase compared to the relative corridor distances one to three and compared to the possible maze group. For this evaluation we can refer to the results described in section “Are the possible and impossible maze comparable?” and interpret them further. For ease of reading we will repeat the description of the analysis here.

First, we performed a 2-way mixed ANOVA on the standard deviation of individual pointing bias. There was a main effect of *target number, F*(1.91, 64.89) = 10.51, *p* < .001, *η_p_^2^* = .24, but no main effect of *maze type*, *F*(1, 34) = 0.00, *p* = .994, *η_p_^2^* < .01, or an interaction, *F*(1.91, 64.89) = 0.09, *p* = .904, *η_p_^2^* < .01. Thus, for both maze groups variability in performance increased with target number. Post-hoc t-tests yielded significant differences between targets one and three, *t*(102) = -3.62, *p* = .003, one and four, *t*(102) = -5.15, *p* ≤ .001, and two and four, *t*(102) = -3.77, *p* = .002, remaining *t*s ≤ 2.23, *p*s ≥ .120. Importantly, no particular increase of noise for distance four in the impossible maze group can be detected. Noise was comparable to the possible maze group and followed a linear pattern across the four targets.

Second, we ran a 2-way ANOVA on pointing latency. Only a main effect of *target number* was found, *F*(1.43, 48.59) = 23.68, *p* < .001, *η_p_^2^* = .41, but no main effect of or interaction with *maze type*, *F*s ≤ 0.66, *p*s ≥ .422. In both mazes, pointing to target one took significantly longer than pointing to the succeeding targets, *t*s ≥ 6.11, *p*s ≤ .001, and differences between targets two to four were not significant, *t*s ≤ 1.09, *p*s ≥ .698. Hence, no particular increase in pointing latency for target four in the impossible maze group can be detected.

In sum, both approaches suggest that participants do not change their procedure to estimate survey relations when facing the fourth target compared to the previous targets within a block. Besides error accumulation across target numbers we failed to find a spike in the withinsubject SD that would suggest greater variability for the impossible maze group in pointing to target four compared to the possible maze group. Likewise, the pointing latency remains constant across target number two, three and four. Hence, participants seem to be rather constant in their estimation behavior. This speaks against the potential strategy that a participant alternates between the biased and the unbiased direction (cw or ccw).

#### 2. Is there a bimodal distribution between subjects?

Besides intra-individual differences in survey estimates also between-subject variation in responding to the fourth target can lead to the examined flattening of the error pattern over relative corridor distance. Figure S1 shows a more detailed version of Figure 5b, where the distribution of the average performance of each participant across all distances is depicted as well. For the impossible maze group the range of responses is highest for relative corridor distance four. The standard deviation of the averaged values across distances are: *SD*_1_ = 13.56, *SD*_2_ = 10.93, *SD*_3_ =15.25, *SD*_4_ = 25.37. While a few participants show a very strong outward bias one participant even shows an inward bias. Hartigans’ dip test for unimodality/multimodality, however, did not yield indications for bimodality or multimodality though, not for the distribution of error pattern for distance four nor the other three distances, *D*s ≤ 0.08, *p*s ≥ .376. Hence, the dip at target four cannot be explained by differences between subjects where some participants followed the biased and others the unbiased sequence for target four. Likewise, we cannot detect groups of different responders, one that points along the local graph prediction, the other that show adjustments towards global embedding. The increase in the between-subject SD seems to be due to a few participants who showed very large outward or inward biases.

**Figure S1.**
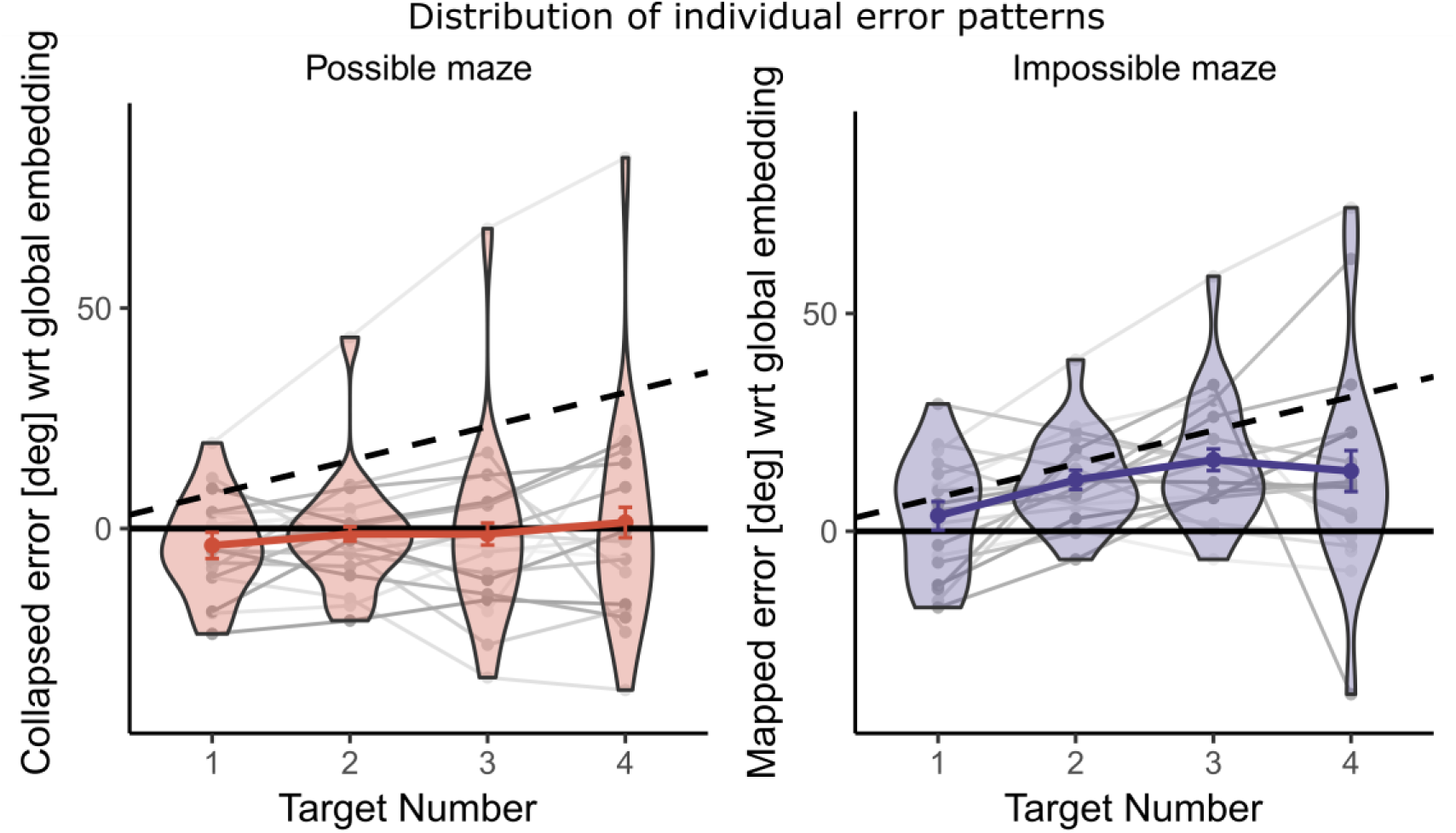
Detailed version of Figure 5b including (in shades of grey) individuals mean performance across target numbers and the distribution of participants average pointing patterns for each corridor distance in the form of a violin plot. In the impossible maze group, the highest standard deviation of inter-individual error variability can be found for relative corridor distance four.

#### 3. Averaging of cw and ccw vector?

Although the analysis of latency and individual standard deviation does not indicate that there is a general change in the way a participant estimates survey relations for the fourth target, we also considered alternative pointing patterns than the ones predicted by the labeled graph and the Euclidean map. One possibility is that participants on average switch from estimating along the biased sequence to estimating along the unbiased direction (e.g., being biased in cw target sequence but recalling target four along the ccw target sequence). If this would be the case a strong inward bias would be expected, which is clearly not reflected in the data.

Alternatively, albeit being biased along one test sequence vectors along both cw and ccw test sequence might have been estimated for target four and then averaged to make a response. In this case a weak outward bias of 6.21° would be predicted for distance four, which is visualized as a violet triangle in Figure 5b^2^. The error pattern of the impossible maze group at relative corridor distance four is not significantly different from this prediction, *t*(17) = 1.27, *p* = .223. Yet, running a Bayesian analysis on the difference of the predicted and the shown error pattern only revealed a weak support for the Null-hypothesis (BF_01_ = 2.07, anecdotal evidence according to Jeffreys, 1961)). Hence, it remains possible that averaging occurred on some trials, which would account for the flattening of the pointing bias to target four.

It should be noted that when running the slope analysis (linear regression on participants individual data points) again while excluding relative corridor distance four the average slope increases to 5.75° (compare to Figure 5c) and is not anymore significantly different from the 7.71° slope predicted for local graph representations, t(17) = -1.65, p = .118. But running a Bayes analysis reveals only anecdotal evidence for the Null-hypothesis, BF_01_ = 1.33, namely, that there is no difference between the calculated slope and the expected slope for the labelled graph model.

In sum our analysis remains to be somewhat inconclusive of what exactly is happening at distance four. It remains that our results revert to a spatial representation in the form of a labeled graph structure and cannot be explained by global embedding into a Euclidean map.

## Footnotes

1 After collapsing the data to represent outwards and inward biases rather than left and rightward biases, we compared the error pattern in the possible and impossible mazes to see whether the pattern of outward biases differed between cw and ccw sequences by performing an ANOVA with the factors *target number* and *test sequence* for each maze group separately. For the impossible maze group, no main effect of *test sequence* and no significant interaction with *target number* was found, *p*’s > .290, indicating comparable error patterns for cw and ccw trials. It was thus valid to merge the collapsed cw and ccw data from the impossible maze. For the possible maze group a significant interaction (*target number x test sequence)* was found, *F*(3, 51) = 3.27, *p* = .029, *η_p_^2^* = .16. This reflects a slight, overall leftward bias in the possible maze group (Figure 4b), which corresponds to a slight outward bias on cw trials and an inward bias on ccw trials. Nevertheless, we decided to merge the collapsed cw and ccw data, given that the leftward bias is small and the possible maze merely provides a baseline.

2 Note that vector averaging for target one, two and three would have led to inward biases, which we clearly did not observe.

## References

Beals, R., Krantz, D. H., & Tversky, A. (1968). Foundations of multidimensional scaling. Psychological Review, 75(2), 127–142. 10.1037/h0025470

Burroughs, W. J., & Sadalla, E. K. (1979). Asymmetries in Distance Cognition. Geographical Analysis, 11(4), 414–421. 10.1111/j.1538-4632.1979.tb00709.x

Byrne, P., Becker, S., & Burgess, N. (2007). Remembering the past and imagining the future: A neural model of spatial memory and imagery. Psychological Review, 114(2), 340–375. 10.1037/0033-295X.114.2.340

Byrne, R. W. (1979). Memory for Urban Geography. Quarterly Journal of Experimental Psychology, 31(1), 147–154. 10.1080/14640747908400714

Chrastil, E. R., & Warren, W. H. (2013). Active and passive spatial learning in human navigation: acquisition of survey knowledge. Journal of Experimental Psychology. Learning, Memory, and Cognition, 39(5), 1520–37. 10.1037/a0032382

Chrastil, E. R., & Warren, W. H. (2014). From Cognitive Maps to Cognitive Graphs. PLoS ONE, 9(11), e112544. 10.1371/journal.pone.0112544

Doeller, C. F., Barry, C., & Burgess, N. (2010). Evidence for grid cells in a human memory network. Nature, 463. 10.1038/nature08704

Ekstrom, a D., Kahana, M. J., Caplan, J. B., Fields, T. a, Isham, E. a, Newman, E. L., & Fried, I. (2003). Cellular networks underlying human spatial navigation. Nature, 425(6954), 184–188. 10.1038/nature01955.1.

Ericson, J. D., & Warren, W. H. (2020). Probing the invariant structure of spatial knowledge: Support for the cognitive graph hypothesis. Cognition, 200, 104276.

Foo, P., Warren, W. H., Duchon, A., & Tarr, M. J. (2005). Do humans integrate routes into a cognitive map? Map-versus landmark-based navigation of novel shortcuts. Journal of Experimental Psychology: Learning, Memory, and Cognition, 31(2), 195–215. 10.1037/0278-7393.31.2.195

Gallistel, C. R. (1990). The Organization of Learning. Cambridge, MA: MIT Press.

Hafting, T., Fyhn, M., Molden, S., Moser, M.-B., & Moser, E. I. (2005). Microstructure of a spatial map in the entorhinal cortex. Nature, 436(7052), 801–806. 10.1038/nature03721

Huttenlocher, J., Hedges, L. V, & Duncan, S. (1991). Categories and particulars: Prototype effects in estimating spatial location. Psychological Review, 98(3), 352–376. Retrieved from http://psycnet.apa.org/journals/rev/98/3/352/

Ishikawa, T., & Montello, D. R. (2006). Spatial knowledge acquisition from direct experience in the environment: Individual differences in the development of metric knowledge and the integration of separately learned places. Cognitive Psychology, 52(2), 93–129. 10.1016/j.cogpsych.2005.08.003

Jacobs, J., Kahana, M. J., Ekstrom, A. D., Mollison, M. V, & Fried, I. (2010). A sense of direction in human entorhinal cortex. Proceedings of the National Academy of Sciences, 107(14), 6487–6492. 10.1073/pnas.0911213107

Jacobs, J., Weidemann, C. T., Miller, J. F., Solway, A., Burke, J. F., Wei, X.-X., … Kahana, M. J. (2013). Direct recordings of grid-like neuronal activity in human spatial navigation. Nature Neuroscience, 16(9), 1188–1190. 10.1038/nn.3466

Jayakumar, R. P., Madhav, M. S., Savelli, F., Blair, H. T., Cowan, J., & Knierim, J. J. (2019). Recalibration of path integration in hippocampal place cells. Nature, 566, 533–537.

Jeffreys, H. (1961). Theory of Probability. Oxford: Oxford University Press

Kluss, T., Marsh, W. E., Zetzsche, C., & Schill, K. (2015). Representation of impossible worlds in the cognitive map. Cognitive Processing, 16(S1), 271–276. 10.1007/s10339-015-0705-x

Kosslyn, S. M., Pick, H. L., & Fariello, G. R. (1974). Cognitive maps in children and men. Child Development, 45(3), 707–716. 10.1111/1467-8624.ep12147012

Loomis, J. M., Klatzky, R. L., Golledge, R. G., Cicinelli, J. G., Pellegrino, J. W., & Fry, P. a. (1993). Nonvisual navigation by blind and sighted: Assessment of path integration ability. Journal of Experimental Psychology: General, 122(1), 73–91. 10.1037/0096-3445.122.1.73

Mallot, H. A., & Basten, K. (2008). Embodied spatial cognition: Biological and artificial systems. Image and Vision Computing, 27(11), 1658–1670. 10.1016/j.imavis.2008.09.001

Marchette, S. A., Ryan, J., & Epstein, R. A. (2017). Schematic representations of local environmental space guide goal-directed navigation. Cognition, 158, 68–80. 10.1016/j.cognition.2016.10.005

McNamara, T. P. (1986). Mental representations of spatial relations. Cognitive Psychology, 18(1), 87–121. 10.1016/0010-0285(86)90016-2

McNamara, T. P., & Diwadkar, V. A. (1997). Symmetry and asymmetry of human spatial memory. Cognitive Psychology, 34(2), 160–190. 10.1006/cogp.1997.0669

McNamara, T. P., Sluzenski, J., & Rump, B. (2008). Human Spatial Memory and Navigation. In H. L. Roediger III (Ed.), Learning and Memory: A Comprehensive Reference (Vol. 2, pp. 157–178). Oxford: Elsevier. 10.1016/B978-012370509-9.00176-5

Meilinger, T. (2008). The Network of Reference Frames Theory: A Synthesis of Graphs and Cognitive Maps. In Spatial Cognition VI. Learning, Reasoning, and Talking about Space (Vol. 5248 LNAI, pp. 344–360). Berlin, Heidelberg: Springer Berlin Heidelberg. 10.1007/978-3-540-87601-4_25

Meilinger, T., Henson, A., Rebane, J., Bülthoff, H. H., & Mallot, H. A. (2018). Humans Construct Survey Estimates on the Fly from a Compartmentalised Representation of the Navigated Environment. In S. Creem-Regehr, J. Schöning, & A. Klippel (Eds.), Spatial Cognition XI. 11th International Conference, Spatial Cognition 2018. Lecture Notes in Artificial Intelligence. Springer.

Meilinger, T., Riecke, B. E., & Bülthoff, H. H. (2013). Local and Global Reference Frames for Environmental Spaces. Quarterly Journal of Experimental Psychology, 67(3), 542–569. 10.1080/17470218.2013.821145

Meilinger, T., Strickrodt, M., & Bülthoff, H. H. (2016). Qualitative differences in memory for vista and environmental spaces are caused by opaque borders, not movement or successive presentation. Cognition, 155. 10.1016/j.cognition.2016.06.003

Meilinger, T., Strickrodt, M., & Bülthoff, H. H. (2018). Spatial Survey Estimation Is Incremental and Relies on Directed Memory Structures. In S. Creem-Regehr, J. Schöning, & A. Klippel (Eds.), Spatial Cognition XI. 11th International Conference, Spatial Cognition 2018. Lecture Notes in Artificial Intelligence (Vol. 11034). Springer.

Moar, I., & Bower, G. H. (1983). Inconsistency in spatial knowledge. Memory & Cognition, 11(2), 107–113. 10.3758/BF03213464

Moeser, S. D. (1988). Cognitive Mapping in a Complex Building. Environment and Behavior, 20(1), 21–49. 10.1177/0013916588201002

Mou, W., McNamara, T. P., Valiquette, C. M., & Rump, B. (2004). Allocentric and egocentric updating of spatial memories. Journal of Experimental Psychology. Learning, Memory, and Cognition, 30(1), 142–157. 10.1037/0278-7393.30.1.142

Muryy, A., & Glennerster, A. (2018). Pointing Errors in Non-Metric Virtual Environments. In S. Creem-Regehr, J. Schöning, & A. Klippel (Eds.), Spatial Cognition XI. 11th International Conference, Spatial Cognition 2018. Lecture Notes in Artificial Intelligence. 10.1101/273532

Newcombe, N., Huttenlocher, J., Sandberg, E., Lie, E., & Johnson, S. (1999). What do misestimations and asymmetries in spatial judgement indicate about spatial representation? Journal of Experimental Psychology: Learning, Memory, and Cognition, 25(4), 986–996. 10.1037/0278-7393.25.4.986

Newcombe, N., & Liben, L. S. (1982). Barrier effects in the cognitive maps of children and adults. Journal of Experimental Child Psychology, 34(1), 46–58. 10.1016/0022-0965(82)90030-3

O’Keefe, J., & Nadel, L. (1978). The hippocampus as a cognitive map. Oxford: Oxford University Press.

Sadalla, E. K., Burroughs, W. J., & Staplin, L. J. (1980). Reference points in spatial cognition. Journal of Experimental Psychology: Humam Learning and Memory, 6(5), 516–528.

Schinazi, V. R., Nardi, D., Newcombe, N. S., Shipley, T. F., & Epstein, R. A. (2013). Hippocampal size predicts rapid learning of a cognitive map in humans. Hippocampus, 23(6), 515–528. 10.1002/hipo.22111

Siegel, A. W., & White, S. H. (1975). The Development of Spatial Representations of Large-Scale Environments. Advances in Child Development and Behavior, 10, 9–55. 10.1016/S0065-2407(08)60007-5

Stern, E., & Leiser, D. (2010). Levels of Spatial Knowledge and Urban Travel Modeling. Geographical Analysis, 20(2), 140–155. 10.1111/j.1538-4632.1988.tb00172.x

Strickrodt, M., Bülthoff, H. H., & Meilinger, T. (2018). Memory for Navigable Space is Flexible and Not Restricted to Exclusive Local or Global Memory Units. Journal of Experimental Psychology: Learning , Memory , and Cognition.

Suma, E. A., Lipps, Z., Finkelstein, S., Krum, D. M., & Bolas, M. (2012). Impossible Spaces: Maximizing Natural Walking in Virtual Environments with Self-Overlapping Architecture. IEEE Transactions on Visualization and Computer Graphics, 18(4), 555–564. Retrieved from http://people.ict.usc.edu/∼suma/papers/suma-vr2012.pdf

Tcheang, L., Bulthoff, H. H., & Burgess, N. (2011). Visual influence on path integration in darkness indicates a multimodal representation of large-scale space. Proceedings of the National Academy of Sciences, 108(3), 1152–1157. 10.1073/pnas.1011843108

Tlauka, M., Carter, P., Mahlberg, T., & Wilson, P. N. (2011). The first-perspective alignment effect: The role of environmental complexity and familiarity with surroundings. The Quarterly Journal of Experimental Psychology, 64(11), 2236–2250. 10.1080/17470218.2011.586710

Tolman, E. C. (1948). Cognitive maps in rats and men. The Psychological Review, 55(4), 189–208. Retrieved from http://psychclassics.yorku.ca/Tolman/Maps/maps.htm

Tversky, B. (1981). Distortions in Memory for Maps. Cognitive Psychology, 13, 407–433. Retrieved from https://www.tc.columbia.edu/faculty/bt2158/faculty-profile/files/1981_Tversky_Distortionsinmemoryformaps.pdf

Wang, R. F. (2016). Building a cognitive map by assembling multiple path integration systems. Psychonomic Bulletin & Review, 23(3), 692–702. 10.3758/s13423-015-0952-y

Wang, R. F., & Brockmole, J. R. (2003). Human navigation in nested environments. Journal of Experimental Psychology: Learning, Memory, and Cognition, 29(3), 398–404. 10.1037/0278-7393.29.3.398

Warren, W. H., Rothman, D. B., Schnapp, B. H., & Ericson, J. D. (2017). Wormholes in virtual space: From cognitive maps to cognitive graphs. Cognition, 166, 152–163. 10.1016/j.cognition.2017.05.020

Weisberg, S. M., Schinazi, V. R., Newcombe, N. S., Shipley, T. F., & Epstein, R. a. (2014). Variations in cognitive maps: Understanding individual differences in navigation. Journal of Experimental Psychology: Learning, Memory, and Cognition, 40(3), 669–682. 10.1037/a0035261

Wilson, P. N., & Wildbur, D. J. (2004). First-perspective alignment effects in a computer-simulated environment. British Journal of Psychology, 95, 197–217. 10.1348/000712604773952421

Wilson, P. N., Wilson, D. a, Griffiths, L., & Fox, S. (2007). First-perspective spatial alignment effects from real-world exploration. Memory & Cognition, 35(6), 1432–44. 10.3758/BF03193613

Zhao, M., & Warren, W. H. (2015a). Environmental stability modulates the role of path integration in human navigation. Cognition, 142, 96–109. 10.1016/j.cognition.2015.05.008

Zhao, M., & Warren, W. H. (2015b). How You Get There From Here. Psychological Science, 26(6), 915–924. 10.1177/0956797615574952

